# Structural determination of human nucleosomes reconstituted by the ExACT platform

**DOI:** 10.64898/2026.07.30.741638

**Authors:** Kei-ichi Okimune, Taiki Azuma, Petra Banko, Junko Haga, Genki Terashi, Ryo Morishita, Yaeta Endo, Shunsuke Kita, Daisuke Kihara, Katsumi Maenaka, Taichi E. Takasuka

**Author notes:** These authors contributed equally: Kei-ichi Okimune, Taiki Azuma. Address correspondence to Taichi E. Takasuka.

## Abstract

The structure and function of eukaryotic chromatin have been extensively studied using conventional salt dialysis-based nucleosome assembly, which has provided fundamental structural insights into nucleosomes as well as a mechanistic understanding of their roles in DNA replication, repair and gene expression. Recently, we developed a labor-saving and time-efficient nucleosome assembly method using a wheat germ cell-free Expression and Assembly Coupled Technology (ExACT), which is emerging as a powerful tool for chromatin research. Here we report cryo-electron microscopy (cryo-EM) structures of human H3.1- and H3.3-containing nucleosomes assembled using this approach and validated by deep-learning-based amino acid-wise model quality (DAQ) scoring. The structures of H3.1- and H3.3-nucleosomes are nearly identical to previously reported models, confirming the structural fidelity of the method. In addition, we determined the previously unreported structure of the primate-specific H3.X-containing nucleosome. Together, these findings validate the cell-free co-expression nucleosome assembly platform and establish this method as a robust framework for biochemically investigating chromatin dynamics.

## Introduction

In eukaryotic cells, the fundamental repeating unit of chromatin is the nucleosome, which comprises two copies each of the core histones H2A, H2B, H3 and H4 wrapped by ∼147 base pairs (bp) of double-stranded DNA^1–3^. Nucleosomes limit the access of DNA-binding proteins to the underlying DNA sequence, and their positions are dynamically regulated via deposition, disassembly, and repositioning by histone chaperones and chromatin remodelers to control essential nuclear processes, including DNA replication^4–6^, repair^7–9^, and transcription^10,11^. Post-translational modifications (PTMs) of core histones and the incorporation of histone variants into nucleosomes are also closely associated with dynamic chromatin organization^12^.

Structural analysis of nucleosomes began soon after chromatin was defined in the 1970s^13^. An initial 20 Å structure of the nucleosome core particle, derived from rat liver nuclei, was limited by sample heterogeneity, including linker histone H1 contamination and nucleosome instability^14^. Subsequent studies achieved a 7 Å resolution, revealing direct interactions between bent DNA and core histones and defining the dyad axis of the nucleosome^15^. High-resolution insight into histone side chains and the DNA path was later obtained using salt dialysis reconstitution with recombinant *Xenopus laevis* histones and symmetric human α-satellite DNA^2^. In 2.8 Å and 1.9 Å crystal structures, over 80% of the histone chain was visualized, revealing the detailed organization of core histone-histone interactions, the superhelical structure of DNA around core histones, and the preference for GC pairs when the major groove faces out, which is offset by ∼5 bp by A-T pairs in the inward minor groove of the DNA double helix^2,16–20^. Meanwhile, the identification of the high-affinity Widom 601 positioning sequence by SELEX further facilitated precise translational positioning of histone octamers for structural studies^21,22^.

Salt dialysis with the Widom 601 sequence has since become a standard method for assembling chromatin in vitro^23–25^. More recently, advances in cryo-electron microscopy (cryo-EM) have enabled the structural determination of diverse nucleosome states and their complexes with regulatory factors, providing profound mechanistic insights into dynamic chromatin organization^26–30^. However, the conventional salt dialysis method remains labor-intensive and time-consuming because it requires multiple steps to prepare the histone octamer prior to the dialysis reaction. Histone octamers are produced either by refolding the denatured histones that have been individually expressed and purified^31,32^, or by co-expressing histones from a polycistronic construct^33^. In either case, additional steps, including the removal of purification tags from histones and chromatographic purification of the histone octamers, are required, and the entire process takes from a week to a month from histone purification to nucleosome reconstitution. Moreover, this non-equilibrium reconstitution bypasses physiological assembly pathways, thereby preventing it from capturing intermediate reactions during chromatin formation in vivo.

To address these drawbacks, we recently reported a wheat germ cell-free Expression and Assembly Coupled Technology (ExACT), where co-expression of the four core histones in the presence of template DNA enables simultaneous protein expression and nucleosome assembly within 20 hours^34–36^. Importantly, the nucleosomes assembled by this method show no apparent post-translational modifications on the histone proteins^37^. As described previously, this approach offers unique features that distinguish it from the conventional salt dialysis method; notably, it enables the capture of chromatin dynamics with non-histone proteins during assembly, which is not feasible with salt dialysis. The wheat cell-free ExACT platform has therefore emerged as a potentially preferred tool for the biochemical assessment of chromatin function in vitro^38^. However, a major limitation of this approach has been the lack of structural information, leaving it elusive whether the assembled nucleosomes are structurally identical to previously reported models.

In this study, we determined the three-dimensional structures of human nucleosomes assembled by the wheat cell-free ExACT platform, including canonical histone H3.1- and histone variant H3.3-containing nucleosomes on the Widom 601 sequence, using cryo-EM. The resulting structures were assessed and compared with previously reported models using the deep-learning-based amino acid-wise model quality (DAQ) score^39^. Furthermore, we demonstrated successful nucleosome assembly using the primate-specific histone variant H3.X, which is evolutionarily derived from H3.3^40^. Although H3.X has been studied in vivo regarding its cell-type-specific expression and associated transcriptional networks^40,41^, structural information for the H3.X-containing nucleosome has been undetermined.

## Results

### Cryo-EM structures of nucleosomes reconstituted via wheat germ cell-free Expression and Assembly Coupled Technology (ExACT)

mRNAs encoding human histones H2A, H2B, H3.1 and H4, together with the histone chaperone Nap1L1, were co-expressed with a 558-bp DNA fragment containing three tandem Widom 601 sequences^21,22,42^ by the wheat cell-free ExACT method (Fig. 1)^34,35^. The reconstituted H3.1-containing nucleosomes (H3.1-nucleosomes) were purified by magnesium-induced precipitation followed by density-gradient ultracentrifugation coupled to chemical crosslinking (GraFix) ^43^. SDS-PAGE analysis of the purified sample confirmed near-stoichiometric levels of all four core histones at the expected molecular weights (Fig. 2a). Electrophoretic mobility shift assay (EMSA) of the same sample yielded a single shifted band corresponding to the assembled nucleosomes (Fig. 2b), indicating homogeneous assembly.

**Fig. 1.**
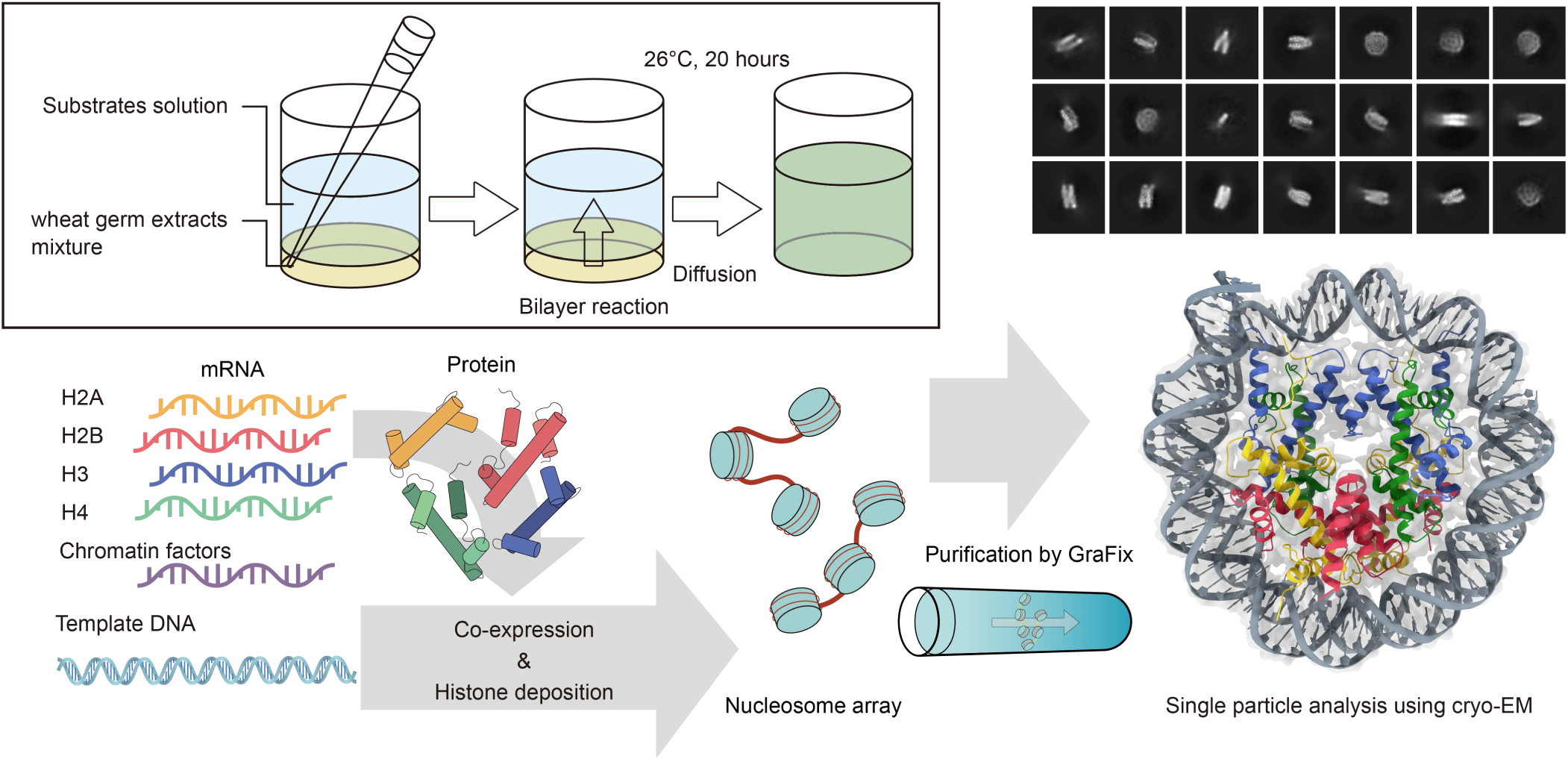
Schematic experimental procedures from co-expression chromatin assembly to cryo-EM structure determination. The ExACT nucleosome reconstitution was initiated as a bilayer reaction in the presence of mRNAs encoding the four core histones and the histone chaperone Nap1L1, alongside a DNA template containing three tandem Widom 601 sequences. The assembled nucleosomal arrays were purified by sucrose gradient ultracentrifugation coupled with chemical crosslinking (GraFix) and subjected to single-particle cryo-EM analysis, which resolved individual mono-nucleosome core particles due to linker flexibility.

**Fig. 2.**
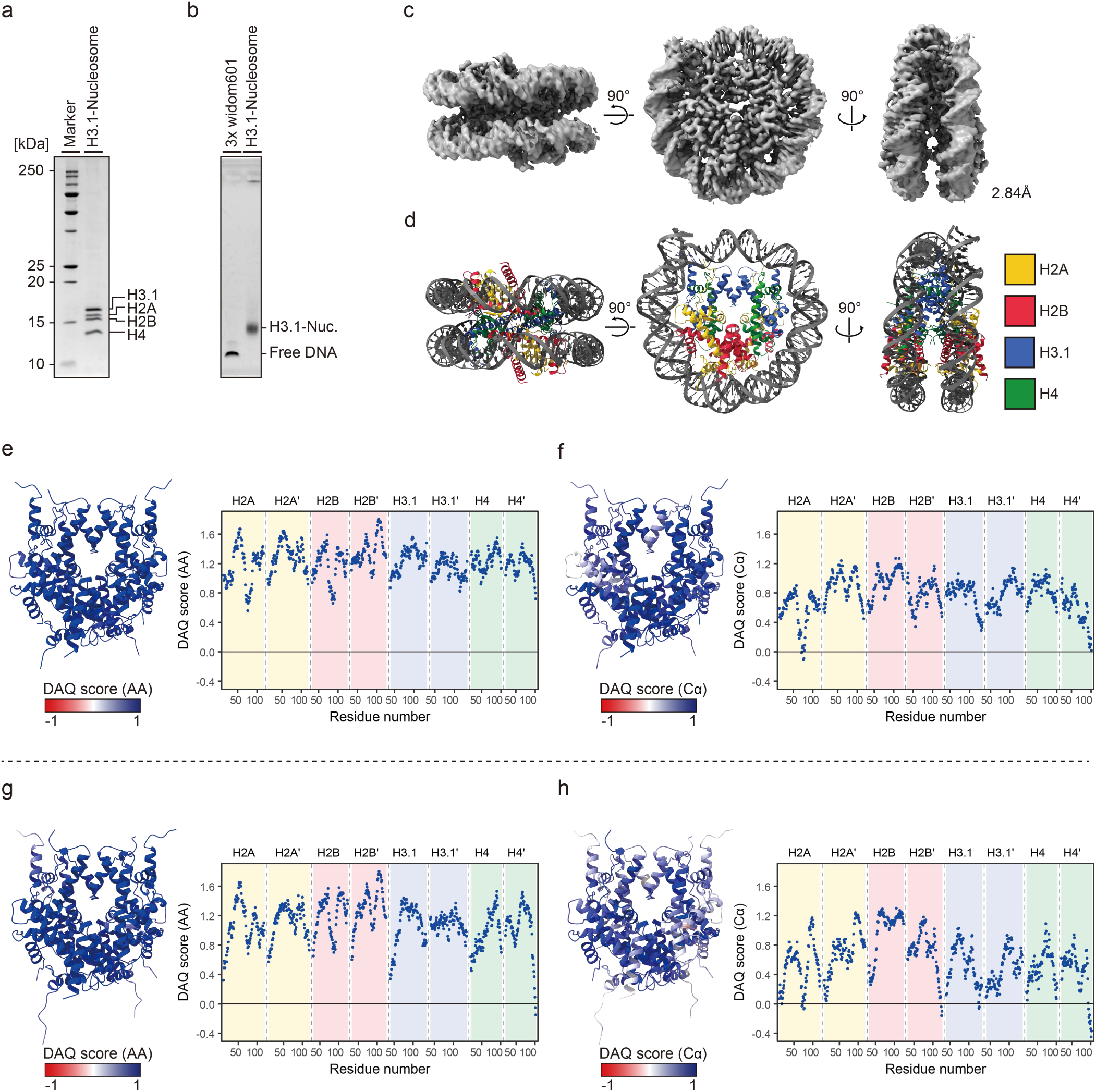
Biochemical and structural verification of assembled H3.1-nucleosomes. **a,** SDS-PAGE analysis of purified H3.1-nucleosomes. **b,** EMSA analysis of purified H3.1-nucleosomes. **c,** 3D electron density map of H3.1-nucleosomes at 2.84 Å resolution. **d,** Cryo-EM structural model of H3.1-nucleosomes. **e**, **f**, Histone octamer of the H3.1-nucleosome colored according to side-chain-wise (**e**) and backbone-wise (**f**) DAQ scores. The color code indicates DAQ values ranging from −1 (poor) to 1 (excellent); the accompanying dot plots show the estimated DAQ score at each residue. **g**, **h**, Cross-validation of our 3D density map against a reported structural model (PDB: 8YBJ) using side-chain-wise (**g**) and backbone-wise (**h**) DAQ scores.

Cryo-EM single particle analysis was performed using 14,809 micrographs processed in cryoSPARC^44^. A total of 347,676 particles were used for 3D reconstruction, yielding a map at 2.84 Å resolution from which an atomic model was built (Fig. 2c,d). The 3D model was validated using the DAQ score^39^ and MolProbity^45^ as shown in Fig. 2e,f (Supplementary Fig. 1 and Supplementary Table 2). The DAQ (AA) and DAQ (Cα) scores were predominantly positive for residues within the core histones, indicating strong local agreement of the amino acid assignments and Cα atom positions with the cryo-EM density map.

Although a zig-zag configuration for tri- and tetra-nucleosomes on the tandem Widom 601 sequence was reported in previous studies using the salt dialysis method, we were unable to reconstruct such tri-nucleosome arrays for the ExACT-assembled H3.1-containing samples^24,28^. We therefore assessed whether the zig-zag conformation could be reconstituted by introducing MgCl₂ at concentrations matching those used in the previous reports during grid preparations, followed by cryo-EM single particle analysis (Supplementary Figs. 2 and 3; Supplementary Table 2). However, the reconstructed density map was indistinguishable from that of the magnesium-free H3.1-mononucleosome, and no ordered tri-nucleosome structure was detected.

### The ExACT-assembled H3.1-nucleosome structure is structurally identical to the reported model

To obtain further structural insights, we cross-validated our 3D density map of the H3.1-nucleosome structure against the reported structure reconstituted by the salt dialysis method (PDB: 8YBJ) (Fig. 2g,h)^46^. The deposited atomic model was fitted into our density map, and the local quality of each residue was evaluated using the DAQ score, encompassing the amino acid assignments and modeled Cα positions. For structural comparison, all side chains in the reported model were precisely refitted into our H3.1-nucleosome density map (Fig. 2g). Although DAQ scores were predominantly positive, a small number of backbone residues in the H2A chain exhibited slightly negative values (Fig. 2h). Overall, these analyses indicate minimal discrepancy between the ExACT-assembled H3.1-nucleosome map and the previously reported structure.

### Structural determination of histone variant H3.3-containing nucleosomes

In addition to canonical histone H3.1, histone H3.3 is a well-characterized variant with defined functions in vivo, for which high-resolution structures are also available^47^. H3.3 differs from H3.1 by only five amino acids: Ser31, Ala87, Ile89, Gly90, and Ser96, with residues 87–90 implicated in chaperone-mediated deposition via pathways dependent on HIRA (Fig. 3a)^48,49^. H3.3-nucleosomes were assembled using the ExACT method and validated by SDS-PAGE and EMSA (Fig. 3b,c). After confirming sample quality, cryo-EM grids were prepared, and 9,544 micrographs yielded a 3D reconstruction at 2.87 Å resolution from 208,786 particles (Fig. 3f,g; Supplementary Fig. 4; Supplementary Table 2). The refined models were validated using the DAQ score and MolProbity (Supplementary Fig. 6; Supplementary Table2).

**Fig. 3.**
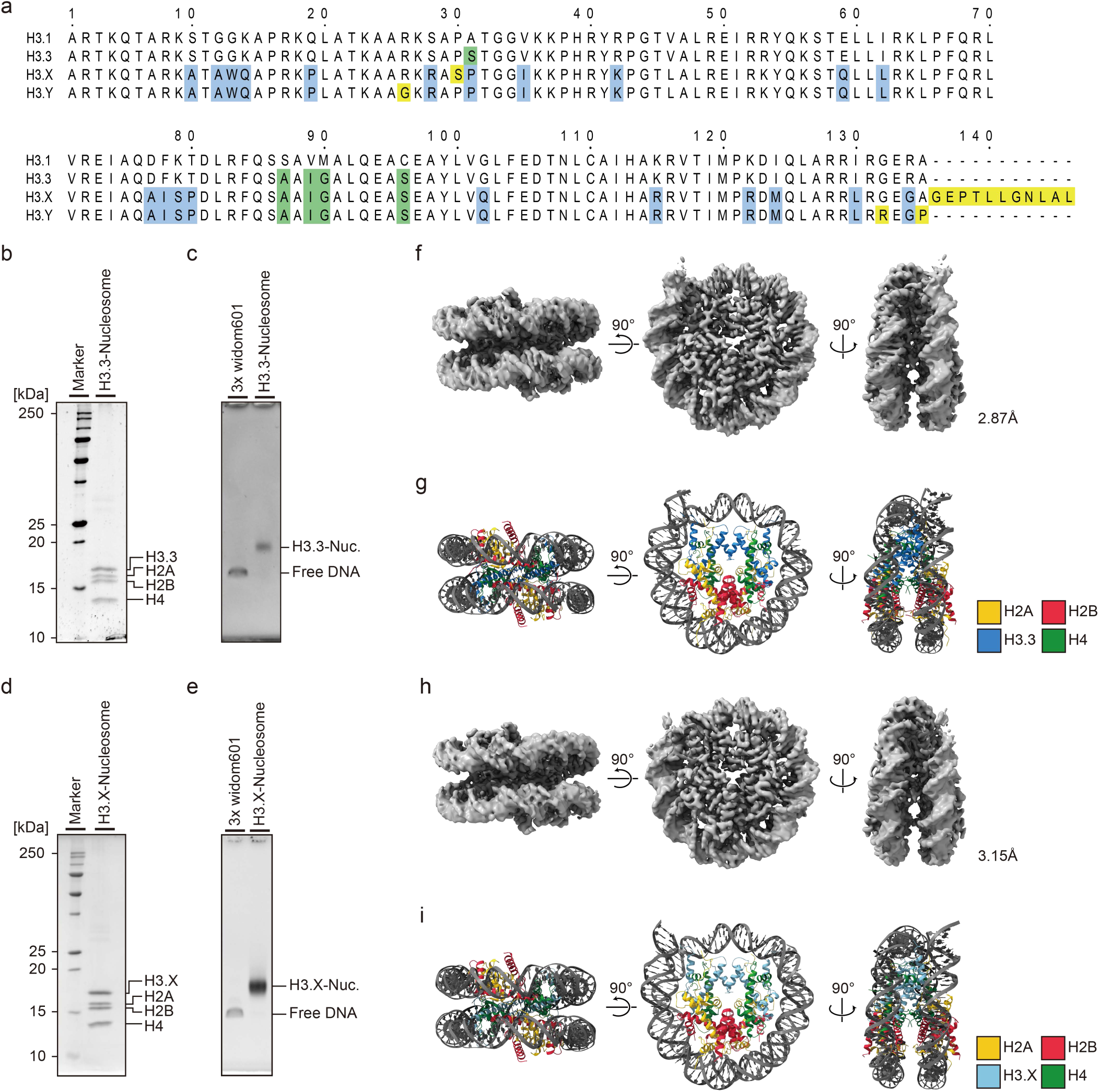
Sample preparation and 3D structure determination of H3.3- and H3.X-nucleosomes. **a**, Multiple sequence alignment of human canonical histone H3.1 and histone variants H3.3, H3.Y, and H3.X. **b–e**, SDS-PAGE (**b**, **d**) and EMSA (**c**, **e**) analyses of reconstituted H3.3 (**b**, **c**) and H3.X (**d**, **e**) nucleosomes. **f**, **g**, 3D electron density map (**f**) and structural model (**g**) of H3.3-nucleosomes at 2.87 Å resolution. **h**, **i**, 3D electron density map (**h**) and structural model (**i**) of H3.X-nucleosomes at 3.14 Å resolution.

Analogous to the structural comparison performed for the H3.1-nucleosome, the H3.3-nucleosomes were cross-validated against the reported structure (PDB: 3AV2)^47^. All side chains in the reported model were precisely refitted into our H3.3-nucleosome density map and DAQ scores were predominantly positive overall (Supplementary Fig. 6), indicating that our ExACT-assembled H3.3-nucleosomes adopt essentially the same structures as those reconstituted by salt dialysis methods.

### The primate-specific histone variant H3.X is efficiently incorporated into nucleosomes

The physiological function of histone H3.X remains poorly understood, and it is only suggested to be a primate-specific histone H3 variant along with its close homolog H3.Y^40^. Both H3.X and H3.Y are thought to be evolutionarily derived from histone H3.3, retaining residues 87-90, but harboring additional distinctive substitutions (Fig. 3a). Specifically, H3.X contains an extended C-terminal tail of 11 amino acids that is absent from H3.3 and H3.Y, and several H3.X and H3.Y-specific amino acid substitutions have been identified (Fig. 3a)^40^. Although the 3D crystal structure of H3.Y-nucleosomes was determined in the previous study using salt dialysis^40^, the structure of the H3.X-nucleosome has remained unreported. We therefore sought to determine the structures of H3.X-nucleosomes to enable direct comparison with the previously reported H3.Y-nucleosome (PDB: 5AY8)^50^ and our H3.3-nucleosome structures.

H3.X-nucleosomes were assembled using the ExACT method and validated by SDS-PAGE and EMSA (Fig. 3d,e). After cryo-EM imaging and data processing, 10,717 micrographs were obtained, and a 3D map was determined at 3.14 Å resolution from 152,207 particles (Fig. 3h,i; Supplementary Fig. 5; Supplementary Table). The resulting model was validated by DAQ score and MolProbity (Supplementary Fig. 6; Supplementary Table 2). The structures between H3.X-nucleosome and the reported H3.3-nucleosome and also with H3.Y-nucleosome were validated by DAQ score (Supplementary Fig. 6e-h). The extended C-terminal region of H3.X was not resolved in the density map (Supplementary Fig. 7c), suggesting conformational flexibility. To assess whether this region had been proteolytically processed during sample preparation process, we performed mass spectrometry on the corresponding SDS-PAGE band (Fig. 3d). LC-MS/MS identified peptides spanning the entire C-terminal region (Supplementary Fig. 7a,b), indicating that the extended tail was intact, hence the C-terminal region is thought to be intrinsically flexible and disordered.

### The DNA entry site of H3.X-nucleosomes exhibit enhanced flexibility compared to H3.3-nucleosomes

To compare H3.3- and H3.X-nucleosomes, we superimposed their structures (Fig. 4). Previous structural and biochemical analyses of H3.3- and H3.Y-nucleosomes highlighted variations at positions Arg42, Lys115, and Lys122 in H3.3 that modulate interactions with the DNA phosphate backbone^50^. In the H3.3-nucleosome, Arg42 engages the DNA phosphate backbone at both entry sites and near the dyad, contributing to tight DNA wrapping (Fig. 4b). In contrast, the substitution of Arg42 with Lys42 in H3.X or H3.Y weakens these interactions near the dyad, resulting in increased flexibility and partial DNA detachment at the entry sites, with an approximate displacement of ∼6.8 Å (Fig. 4a,c).

**Fig. 4.**
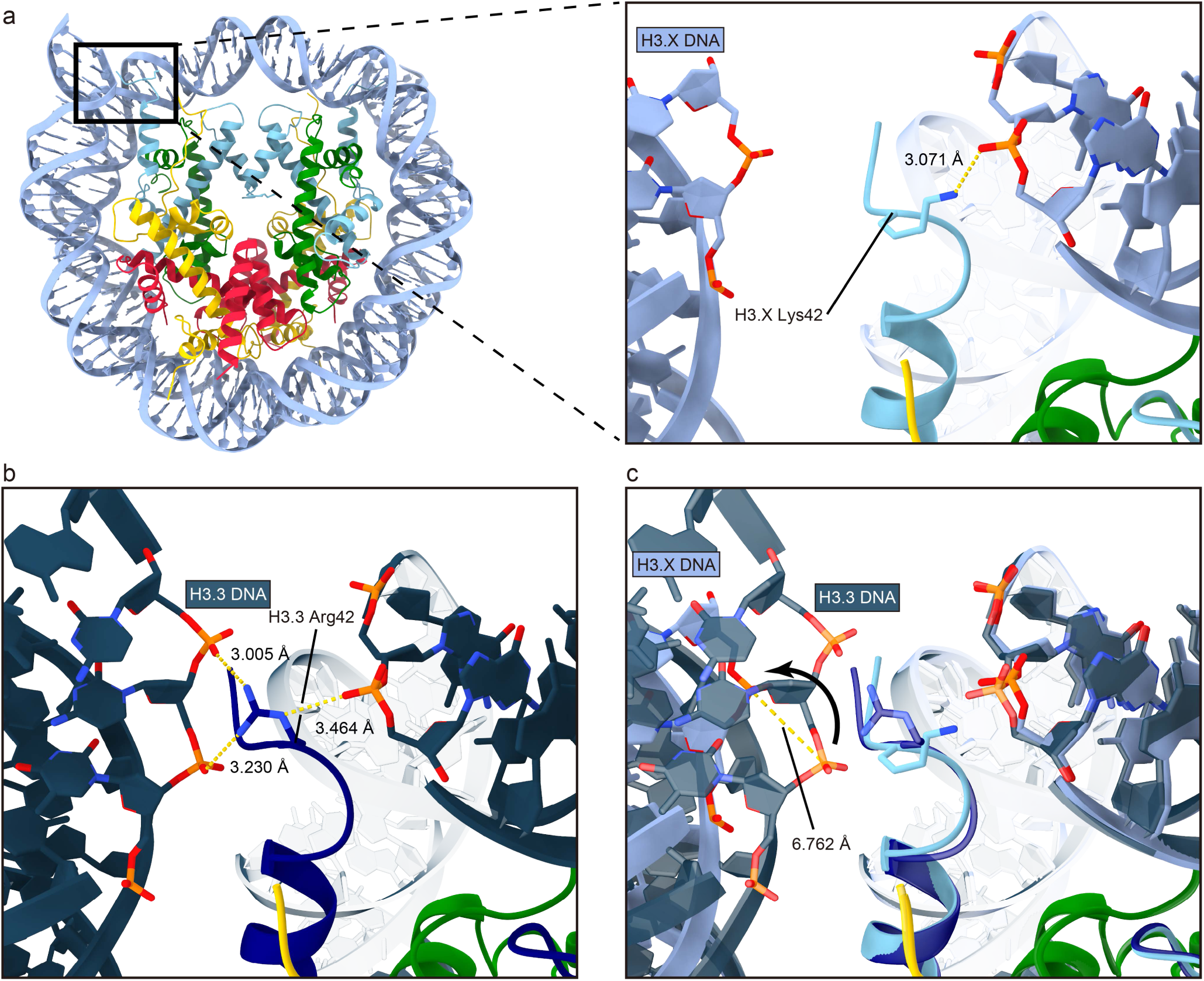
3D structure of H3.X and structural comparison with H3.3. **a**, A front view of H3.X-nucleosome (left) with close-up view of the DNA-histone interaction of Lys42 in H3.X (right) at the DNA entry site. **b**, Close-up views of the DNA-histone interaction of Arg42 in H3.3. **c**, Superimposed view of H3.X- and H3.3-nucleosomes highlighting conformational differences at the DNA entry sites.

Although previous studies suggested the functional importance of these substitutions, such DNA destabilization at the DNA entry site was not observed in earlier X-ray structures (PDB: 5AY8)^50^, possibly due to DNA compaction imposed by crystal packing constraints. Together, these findings indicate that the Arg42Lys substitution destabilizes DNA interactions at the entry sites of H3.X-nucleosomes, potentially influencing downstream DNA-templated processes such as transcription or remodeling.

## Discussion

Our previous work demonstrated that co-expressing the four core histones using the wheat germ cell-free reactions enables efficient nucleosome assembly in vitro^34–37^. However, the resulting products were evaluated only by classical biochemical assays, and a robust method for structural determination was needed. Here, we showed that the combination of magnesium-induced precipitation and GraFix provides a reliable method for isolating the reconstituted nucleosomes with high purity. Our cryo-EM analysis and structural validation by DAQ score demonstrate that our nucleosome structures are virtually identical to the reported models. DAQ (AA) and DAQ (Cα) scores quantify residue- and backbone-wise local model quality from cryo-EM density maps, respectively. The predominantly positive scores across all histone chains indicate close agreement between the H3.1-nucleosomes from the current study and those reported previously. It is also important to note that our cryo-EM analysis of the reconstituted sample did not reveal the previously described zig-zag configuration, indicating that this arrangement is infrequently present under our reconstitution conditions, which aligns with observations in vivo^24,28,51^.

Beyond canonical nucleosomes, we determined cryo-EM structures of nucleosomes containing the histone H3 variants H3.3 or H3.X. While our models showed overall positive correlations with reported H3.3- and H3.Y-nucleosomes by residue-wise DAQ (AA) comparison, their DAQ (Cα) scores indicated local backbone discrepancies (Supplementary FIg.6e-h). This discordance reflects the design of the DAQ framework, in which amino acid identity and backbone position are evaluated as separate density-derived features in cryo-EM structures^52^. In particular, the nucleosome structures determined by two distinct methods, X-ray crystallography versus cryo-EM, might contribute to this result. For example, extensive overlap between protein and DNA density at histone-DNA interfaces can reduce the distinctiveness of backbone-associated features while preserving residue-specific density patterns. Moreover, nucleosomes exhibit conformational heterogeneity arising from DNA breathing, transient DNA unwrapping, and local histone rearrangements, all of which might broaden backbone density in reconstructed cryo-EM maps. Such modeling procedures generally preserve sequence registration but may retain subtle backbone inaccuracies when conformational differences exist between the crystal structures and the cryo-EM structures.

The extended C-terminus of H3.X was not resolved, suggesting conformational flexibility and potential interactions with histone-modifying enzymes, chaperones, or chromatin remodeling factors. Furthermore, the substitution of Arg42 with Lys in H3.X, as compared to H3.3, increased DNA flexibility at the entry site, with potential consequences for DNA-templated processes such as transcription. These structural validations support our wheat cell-free ExACT platform. Moreover, coupling this platform with cryo-EM analysis, as established here, provides a general strategy for determining the structures of more complex assemblies containing non-histone chromatin proteins in addition to nucleosomes. This approach is particularly advantageous for proteins that are insoluble when expressed in *E. coli* and for capturing transient or dynamic intermediates formed during nucleosome assembly.

### Remarks

Cell-free co-expression of core histones provides a one-step reaction that achieves simultaneous histone expression and nucleosome assembly without denaturing and refolding processes, resulting in a highly streamlined and time-efficient reconstitution workflow. Histone co-expression has been accepted as a strategy to simplify the required processes for nucleosome assembly in vitro^33^. Recently, alternative methods that assemble nucleosomes in the *E. coli* cells using polycistronic expression have been engineered^53–56^. This approach shares with our method the feature that histone synthesis and nucleosome assembly are coupled. Although *E. coli* chromatinization has the advantage of assembling nucleosomes in a cellular context, it may be constrained by the difficulty of balancing differences in the expression efficiencies of individual histone proteins and by limitations on the amounts and types of external proteins that can be synthesized to maintain host-cell viability. In contrast, our ExACT platform overcomes differences in translation efficiency by adjusting the abundance of each input mRNA^34^ and allows further protein synthesis independent of cell viability^36^. If our platform can be extended to efficient nucleosome reconstitution on much longer DNA templates, such as genomic DNA, it will provide a stable environment where multiple proteins co-exist functionally, thereby bridging in vitro conditions and the in vivo context^57^.

## Method

### Cloning

Human histones H2A, H2B, H3.1 and H4 were prepared as described previously^34^. Coding sequences for H3.3 (UniProt: P84243), H3.X (UniProt: P0DPK5) and Nap1L1 (UniProt: P55209) were amplified from a human cDNA library using the primer pairs listed in Supplementary Table S1 and cloned into the pEU-E01-LICNot vector via ligation-independent cloning^58^. Constructs were transformed into DH5α competent cells (SMOBiO, Hsinchu, Taiwan). Transformants were screened by colony PCR (Supplementary Table 1) and verified by DNA sequencing (Eurofins, Tokyo, Japan). Plasmids were purified using a NucleoBond Xtra Midi kit (MACHEREY-NAGEL, Dueren, Germany). The tandem Widom 601 sequence was prepared as described previously^35^. Briefly, a gel-purified linear DNA fragment was generated from pBS_12x-601_MA2 (Addgene, Watertown, MA) using *Sml*I (New England Biolabs, Ipswich, MA) digestion and a 558-bp fragment containing the 3xWidom 601 sequence was amplified by PCR using the primer pair listed in Supplementary Table 1.

### In vitro transcription

mRNAs encoding human histones (H2A, H2B, H3.1, H3.3, H3.X, and H4) and the histone chaperone Nap1L1 were synthesized individually by in vitro transcription as described previously ^34^. Briefly, 8.0 μg of template plasmid DNA for each gene was incubated with 8 units of SP6 RNA polymerase (Promega, Madison, WI) in 1x transcription buffer (CellFree Sciences, Yokohama, Japan) for 4 h at 37°C. The synthesized mRNAs were treated with 12-16 unit μl^-1^ DNase I (NIPPON GENE, Tokyo, Japan) for 1 h at 37 °C, deproteinized using a standard protocol, and quantified using a NanoDrop spectrophotometer (Thermo Fisher Scientific, Waltham, MA).

### ExACT reaction

A reaction mixture containing 10 μl of wheat germ extract WEPRO7240Ch (CellFree Sciences), 0.8 μg of creatin kinase (Sigma-Aldrich, St. Lousi, MO), 0.5 μg of 3x Widom 601 DNA, and designated amounts of histone and Nap1L1 mRNAs was adjusted to a total volume of 20.8 μl with nuclease-free water in a 96-well plate. The mixture was underlaid with 206 μl 1× SUB-AMIX (CellFree Science) and incubated for 20 h at 26 °C. For structural analysis, 46 individual reactions were pooled per sample. For the reconstitution of nucleosomes containing H3.1, H3.3, or H3.X, mRNA amounts were optimized as follows: 6 μg each of H2A and H2B mRNAs, 11 μg of H4 mRNA, 7 μg of Nap1L1 mRNA, and either 3 μg of H3.1, 3 μg of H3.3, or 11 μg of H3.X mRNA.

### Purification of nucleosomes

Assembled nucleosomes were purified by magnesium ion-induced precipitation followed by GraFix^43^. Briefly, MgCl₂ was added to the reaction mixture to a final concentration of 10 mM. The mixture was incubated for 15 min at 26 °C and then centrifuged at 12,000 × g for 10 min at 4 °C. The resultant precipitate was washed with a buffer containing 10 mM HEPES–NaOH (pH 7.5), 20 mM NaCl, 1 mM DTT and 10 mM MgCl₂, and centrifuged at 12,000 x g for 5 min at 4 °C. The pellet was resuspended in 10 mM HEPES–NaOH (pH 7.5), 20 mM NaCl, 1 mM DTT and 1 mM EDTA, and incubated for 120 min at 55 °C to thermally position the histone octamer at the center of the Widom 601 sequence. The sample was then applied to the top of a 5–20% sucrose gradient containing 0–4% paraformaldehyde in 10 mM HEPES–NaOH (pH 7.5), 20 mM NaCl and 1 mM DTT, and centrifuged at 26,000 rpm for 16 h at 4 °C in an SW60Ti rotor (Beckman Coulter, Brea, CA, USA). Fractions corresponding to the target tri-nucleosomes were concentrated using a Vivaspin 500 centrifugal concentrator (MWCO 30,000, PES; Sartorius, Gottingen, Germany) and buffer-exchanged three times into 20 mM HEPES-KOH (pH 7.5) and 1 mM DTT to remove residual sucrose and paraformaldehyde.

### Denaturing and native polyacrylamide gel electrophoresis

Purified nucleosomes were analyzed by 18% SDS–PAGE and visualized by Coomassie Brilliant Blue or Oriole fluorescent gel staining. The intact structural integrity of the assemblies was examined by electrophoretic mobility shift assay (EMSA) using 0.8% agarose gels in 0.5× TBE buffer, followed by ethidium bromide staining.

### Cryo-EM data acquisition

A 3 µL aliquot of purified samples (0.05 µg µl^-1^) was applied onto a freshly glow-discharged Quantifoil R1.2/1.3 200-mesh Cu grid (Quantifoil Micro Tools GmbH, Jena, Germany). Grids were blotted for 3 s at 18 °C and 100% humidity using a Vitrobot Mark IV (Thermo Fisher Scientific) and plunge-frozen in liquid ethane. Micrographs were acquired on a Titan Krios G4 transmission electron microscope (Thermo Fisher Scientific) operated at 300 kV and equipped with a K3 direct electron detector and a GIF Bio-Continuum energy filter (Gatan, Pleasanton, CA) utilizing a 20 eV slit width. Data collection was performed at a nominal magnification of ×105,000, resulting in a physical pixel size of 0.85 Å per pixel. For H3.1-nucleosomes, 9,910 and 14,809 movies were collected in the presence or absence of 1 mM MgCl₂, respectively. For H3.3-and H3.X-nucleosomes, 9,544 and 10,717 movies were collected, respectively. All datasets were recorded automatically using EPU software with an exposure time of 1.5 s, yielding a total electron dose of 48∼9 e^-/Å2^ fractionated over 50 frames. The nominal defocus range was set from −0.8 to −2.0 μm.

### Cryo-EM data processing

All datasets were processed independently using cryoSPARC (v4.7.1) (Structura Biotechnology, Toronto, Canada). For H3.1-containing nucleosomes in Mg^2+^-free buffer, movie frames were aligned and dose-weighted via Patch motion correction, and contrast transfer function (CTF) parameters were estimated using Patch CTF estimation. Template 2D classes were generated by blob picking followed by 2D classification from 200 representative micrographs. A total of 9,953,169 particles were template-picked, and two rounds of reference-free 2D classification (200 classes, 40 iterations) were performed to remove poorly defined particles and background artifacts. A subset of 732,470 particles was selected for ab initio reconstruction to generate initial 3D volumes. Heterogeneous refinement was carried out to further partition discrete particle populations. Finally, 347,676 particles were retained for final map reconstruction and refinement using non-uniform refinement with global and local CTF refinement, followed by local refinement.

For H3.1-containing nucleosomes in the presence of 1 mM MgCl₂, movie frames underwent Patch motion correction and Patch CTF estimation. Following initial blob picking and 2D classification from 200 micrographs to generate templates, 5,998,933 particles were template-picked. Reference-free 2D classification (200 classes, 40 iterations) eliminated poor particles, yielding 582,516 particles. Three rounds of ab initio reconstruction and heterogeneous refinement were carried out to eliminate low-resolution structures. Ultimately, 88,640 particles were isolated for final map reconstruction via non-uniform refinement and local refinement.

For H3.3-containing nucleosomes, movie frames were processed using Patch motion correction and Patch CTF estimation. Following template generation from 200 micrographs, 5,359,496 particles were template-picked. Two rounds of reference-free 2D classification (200 classes, 40 iterations) filtered out poorly defined particles, leaving 974,982 particles. Two iterations of ab initio reconstruction and heterogeneous refinement successfully excluded low-resolution structures, thereby isolating 208,786 particles for final map reconstruction using non-uniform and local refinement.

For H3.X-containing nucleosomes, movie frames were aligned and dose-weighted via Patch motion correction, and CTF parameters were determined using Patch CTF estimation. Template 2D classes were generated from 200 micrographs. A total of 7,216,039 particles were template-picked, and two rounds of reference-free 2D classification (200 classes, 40 iterations) were executed to discard low-quality particles. A subset of 435,429 particles was chosen for ab initio reconstruction to yield initial models, followed by heterogeneous refinement to resolve structural heterogeneity. After local motion correction, 152,207 particles were retained for final map reconstruction and non-uniform refinement with global and local CTF refinement, followed by local refinement.

Global resolutions for all final maps were determined based on the gold-standard Fourier shell correlation criterion (FSC = 0.143). Data collection and processing statistics are summarized in Supplementary Tables 2.

### Model building

Model building Initial structural models of H3.1-, H3.3-, and H3.X-containing nucleosomes were generated using the cryo-EM structure of the canonical core histones wrapped by 145 bp of Widom 601 DNA (PDB: 8JBX)^59^ as a starting template. Amino acid sequences of H2A and H3 were manually modified to correspond to human H2A type 2-A and human H3.1, H3.3 or H3.X, respectively. Model coordinates were refined by real-space refinement in Phenix^60^ and manually adjusted in Coot^61^. Structural figures were prepared using ChimeraX^62^ and PyMOL^63^. Final models were validated using MolProbity and DAQ-score metrics^39^.

### Model comparison

For H3.1-containing nucleosomes, a previously reported model (PDB: 8YBJ) was fitted into our experimental density map via rigid-body refinement in Phenix, treating all chains as a single rigid body. Model-to-map validation was evaluated using DAQ-score and MolProbity parameters. For H3.3- and H3.X-containing nucleosomes, structures were superimposed, and the distance of the DNA backbones was calculated using ChimeraX.

### Mass spectrometry analysis of H3.X

Mass spectrometry analysis of H3.X Purified H3.X-containing nucleosomes reconstituted by the cell-free method were analyzed via in-gel digestion LC-MS/MS using a modified protocol^64^. Briefly, 12 μl of purified nucleosome sample was resolved on an 18% acrylamide SDS–PAGE gel and stained with Coomassie Brilliant Blue. The protein band corresponding to H3.X was excised into ∼1 × 1 mm pieces and destained sequentially with 300 μl of 25 mM ammonium bicarbonate in 50% methanol, followed by 300 μl of 25 mM ammonium bicarbonate in 50% acetonitrile for 30 min at 40 °C. Gel pieces were reduced with 50 μl of 10 mM DTT in 25 mM ammonium bicarbonate for 45 min at 56 °C, and subsequently alkylated with 50 μl of 55 mM iodoacetamide in 25 mM ammonium bicarbonate for 30 min at room temperature in the dark. Enzymatic digestion was carried out overnight at 37 °C by adding 1 μl of Arg-C protease (20 ng μl⁻¹; Promega). Tryptic peptides were extracted sequentially using 50 μl of 50% acetonitrile/0.1% formic acid (accompanied by 10 min of sonication), 100% acetonitrile/0.1% formic acid, and 100% H₂O/0.1% formic acid, and purified using ZipTip C18 pipette tips (Millipore).

Proteomic analysis was performed on an Easy-nLC 1000 system coupled to a Q Exactive Plus Orbitrap mass spectrometer (Thermo Fisher Scientific) operating via Xcalibur v3.1 software. Peptides were separated on a C18 capillary column (NTCC-360/75-3-125; Nikkyo Techno, Tokyo, Japan) using a 120-min linear gradient from 0% to 40% mobile phase B (0.1% formic acid in acetonitrile) against mobile phase A (0.1% formic acid in water) at a constant flow rate of 300 nl min ^-1^. Full-scan MS spectra were acquired across a mass range of 300–2,000 m/z at a resolution of 70,000, with a maximum injection time of 50 ms and an AGC target of 3e6. Peptide identification and quantification were performed using Proteome Discoverer v2.1 (Thermo Fisher Scientific) searched against the targeted human H3.X primary sequence. Precursor and fragment mass tolerances were constrained to 10 ppm and 0.6 Da, respectively.

### Data availability

3D density maps and atomic models of each series of nucleosomes were deposited to the PDB and EMDB under accession code EMD-81900 and 43IB (human canonical nucleosomes in the absence of 1mM MgCl_2_), EMD-81902 and 43IE (human canonical nucleosomes in the presence of 1mM MgCl_2_), EMD-81903 and 43IF (human H3.3-containing nucleosomes), and EMD-81904 and 43IG (human H3.X containing nucleosomes). Mass spectrometry analysis data is available via PRIDE database under accession code PXD080480.

## Acknowledgements

This work was partly supported by the Institute for Fermentation, Osaka (IFO) (to T.E.T.), AMED grant for Basis for supporting innovative drug discovery and life science research phase 1 JP17am0101093 (to K.M.) and phase 2 JP22ama121037 (to K.M.). K.O. was supported by JSPS Research Fellowships for Young Scientists DC1 (22KJ0120). J.H. and T.A. were supported by JST SPRING, Grant Number JPMJSP2119. D.K. was partially supported by National Institutes of Health (R35GM158267). G.T. was supported by the National Science Foundation (DBI2433490).

## Author information

### Authors and Affiliations

**Graduate School of Global Food Resources and Research Faculty of Agriculture, Hokkaido University, Sapporo, Japan**

Kei-ichi Okimune, Taiki Azuma, Petra Banko, Junko Haga & Taichi E. Takasuka

**Department of Biological Sciences, Purdue University, West Lafayette, United States of America**

Genki Terashi & Daisuke Kihara

**Department of Computer Science, Purdue University, West Lafayette, United States of America**

Daisuke Kihara

**CellFree Sciences Co., Ltd, Matsuyama, Japan**

Ryo Morishita

**Ehime Prefectural University of Health Sciences, Iyo-gun, Japan**

Yaeta Endo

**Laboratory of Biomolecular Science, and Center for Research and Education on Drug Discovery, Faculty of Pharmaceutical Sciences, Hokkaido University, Sapporo, Japan**

Shunsuke Kita & Katsumi Maenaka

**Division of Pathogen Structure, International Institute for Zoonosis Control, Hokkaido University, Sapporo, Japan and Faculty of Pharmaceutical Sciences, Kyushu University, Fukuoka, Japan**

Katsumi Maenaka

### Contributions

K.O, Y.E., and T.E.T. designed research; K.O., T.A., and R.M. prepared human histones and carried out in vitro wheat germ co-expression chromatin assembly reactions. K.O., T.A., J.H., S.K., G.T., D.K., and K.M. performed cryo-EM experiments, including data acquisition, image processing and built the cryo-EM models. K.O. and T.A. prepared the figures; K.O., T.A., and T.E.T. wrote the paper and revised by all authors.

### Ethics declarations

The authors show no ethics declarations.

### Supplementary information

Supplementary Figs. 1-7 and Supplementary Table 1,2.

### Rights and permissions

Springer Nature or its licensor (e.g. a society or other partner) holds exclusive rights to this article under a publishing agreement with the author(s) or other rightsholder(s); author self-archiving of the accepted manuscript version of this article is solely governed by the terms of such publishing agreement and applicable law.

