## Supplementary information for "Structural determination of human nucleosomes reconstituted by the ExACT platform"

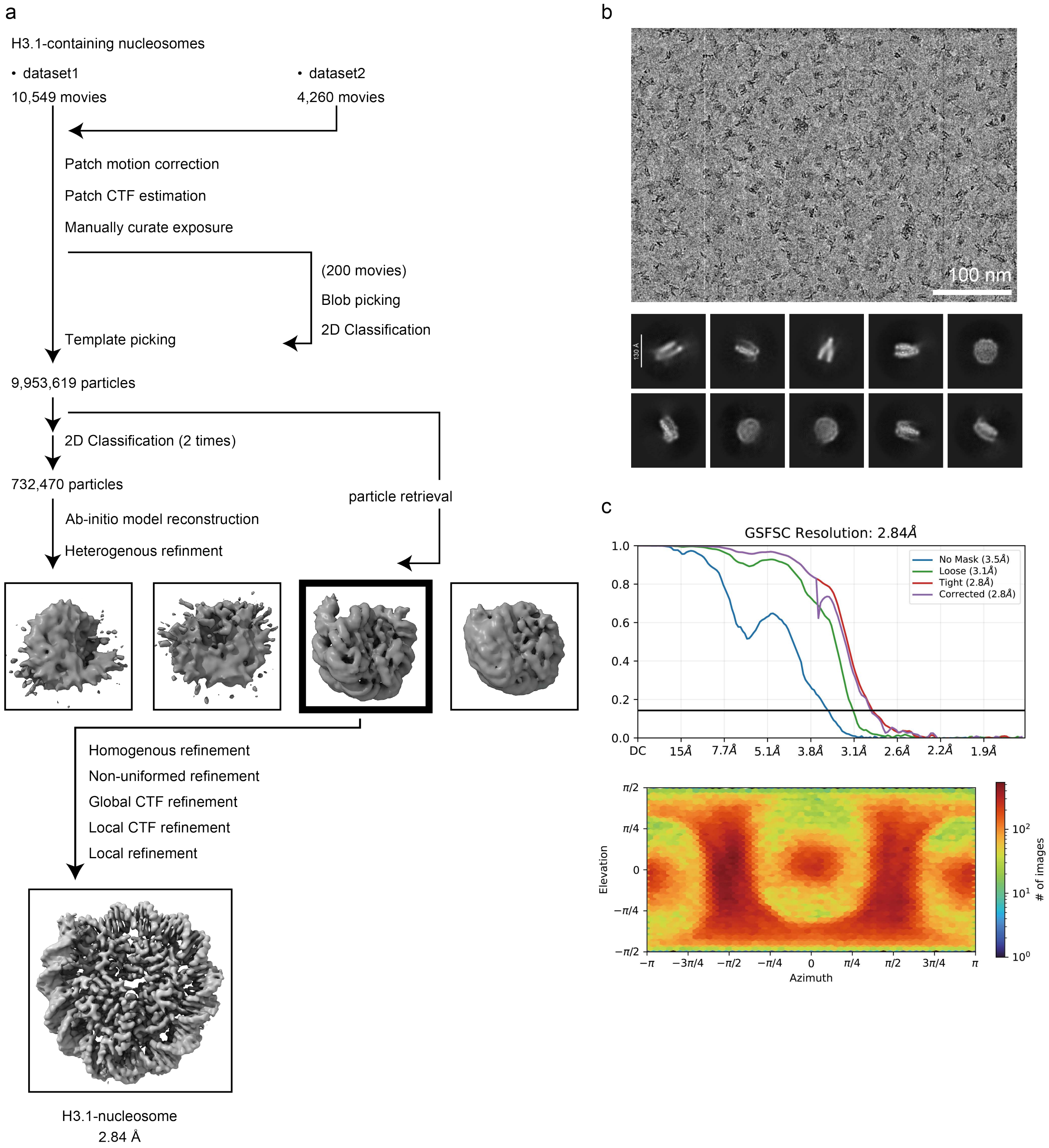


**Supplementary Figure 1. Data processing of H3.1-containing nucleosomes.**

**a,** Cryo-EM data processing workflow for H3.1-containing nucleosomes in the absence of $MgCl_2$. **b,** Representative micrograph and 2D class images. **c,** Global resolution assessment by gold-standard Fourier shell correlation (FSC) curves at the 0.143 criteria (top) and the angular distribution of particle representation by viewing direction distribution plot (bottom).

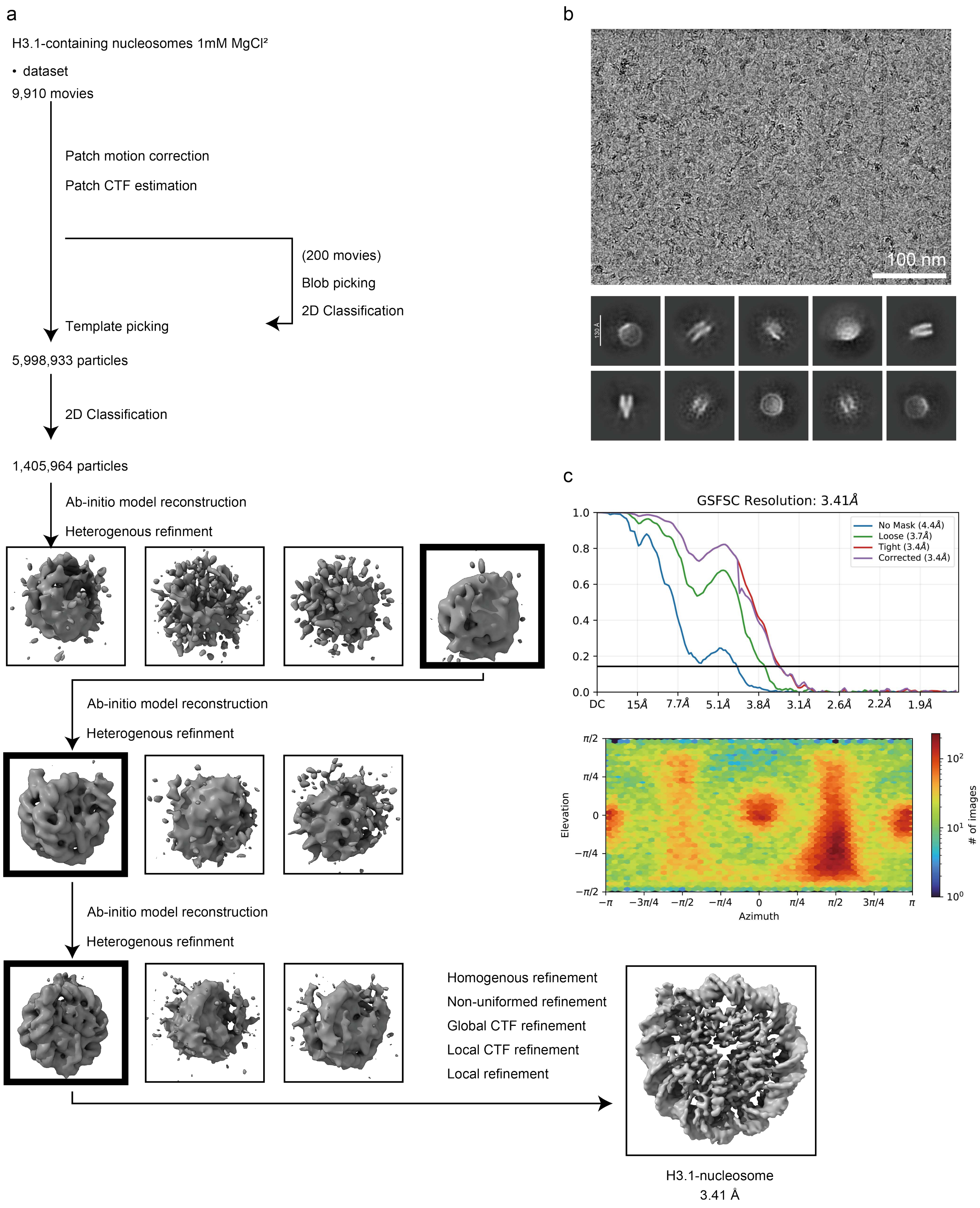


**Supplementary Figure 2. Data processing of H3.1-containing nucleosomes in the presence of $Mg^{2+}$.**
**a**, Cryo-EM data processing workflow for H3.1-containing nucleosomes in the presence of 1 mM $MgCl_2$. **b**, Representative micrograph and 2D class images. **c**, Global resolution assessment by FSC curves at the 0.143 criteria (top) and the angular distribution of particle representation by viewing direction distribution plot (bottom).

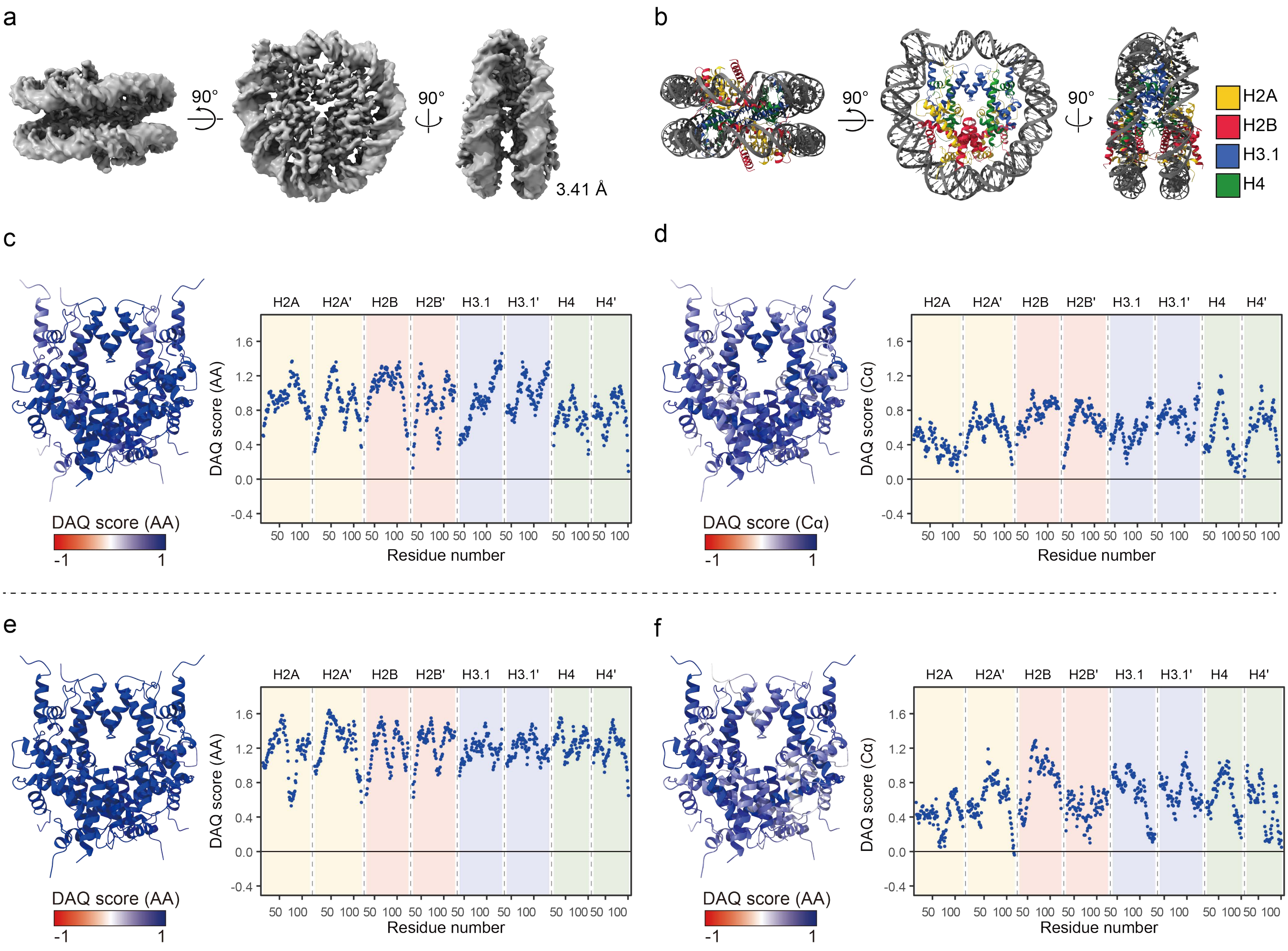


**Supplementary Figure 3. Cryo-EM structures and validation of H3.1-containing nucleosomes determined in the presence of 1 mM $MgCl_2$.**

**a,** 3D electron density map of H3.1-nucleosomes at 3.41 Å resolution in the presence of 1 mM $MgCl_2$. **b,** Cryo-EM structure model of H3.1-nucleosomes in the presence of 1 mM $MgCl_2$. **c,d,** Histone octamer of the H3.1-nucleosome in the presence of 1 mM $MgCl_2$ colored according to side-chain-wise (**c**) and backbone-wise (**d**) DAQ scores. The color code indicates DAQ values ranging from -1 (poor) to 1 (excellent); the accompanying dot plots show the estimated DAQ score at each residue. **e**,**f**, Cross-validation of our 3D density map in the absence of $MgCl_2$ against our structural model in the presence of $MgCl_2$ using side-chain-wise (**e**) and backbone-wise (**f**) DAQ scores.

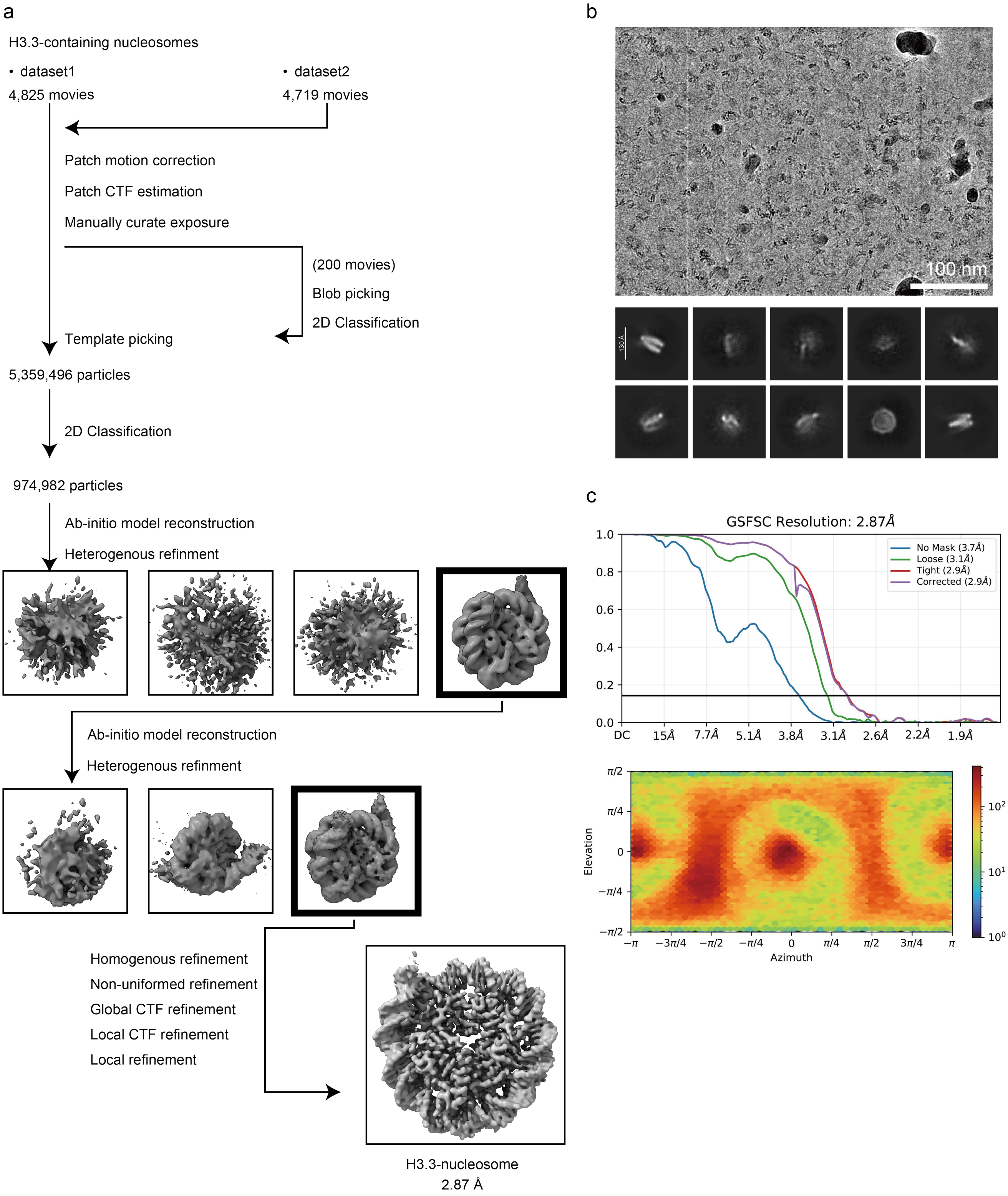


**Supplementary Figure 4. Data processing of H3.3-containing nucleosomes.**

**a,** Cryo-EM data processing workflow for H3.3-containing nucleosomes. **b,** Representative micrograph and 2D class images. **c,** Global resolution assessment by FSC curves at the 0.143 criteria (top) and the angular distribution of particle representation by viewing direction distribution plot (bottom).

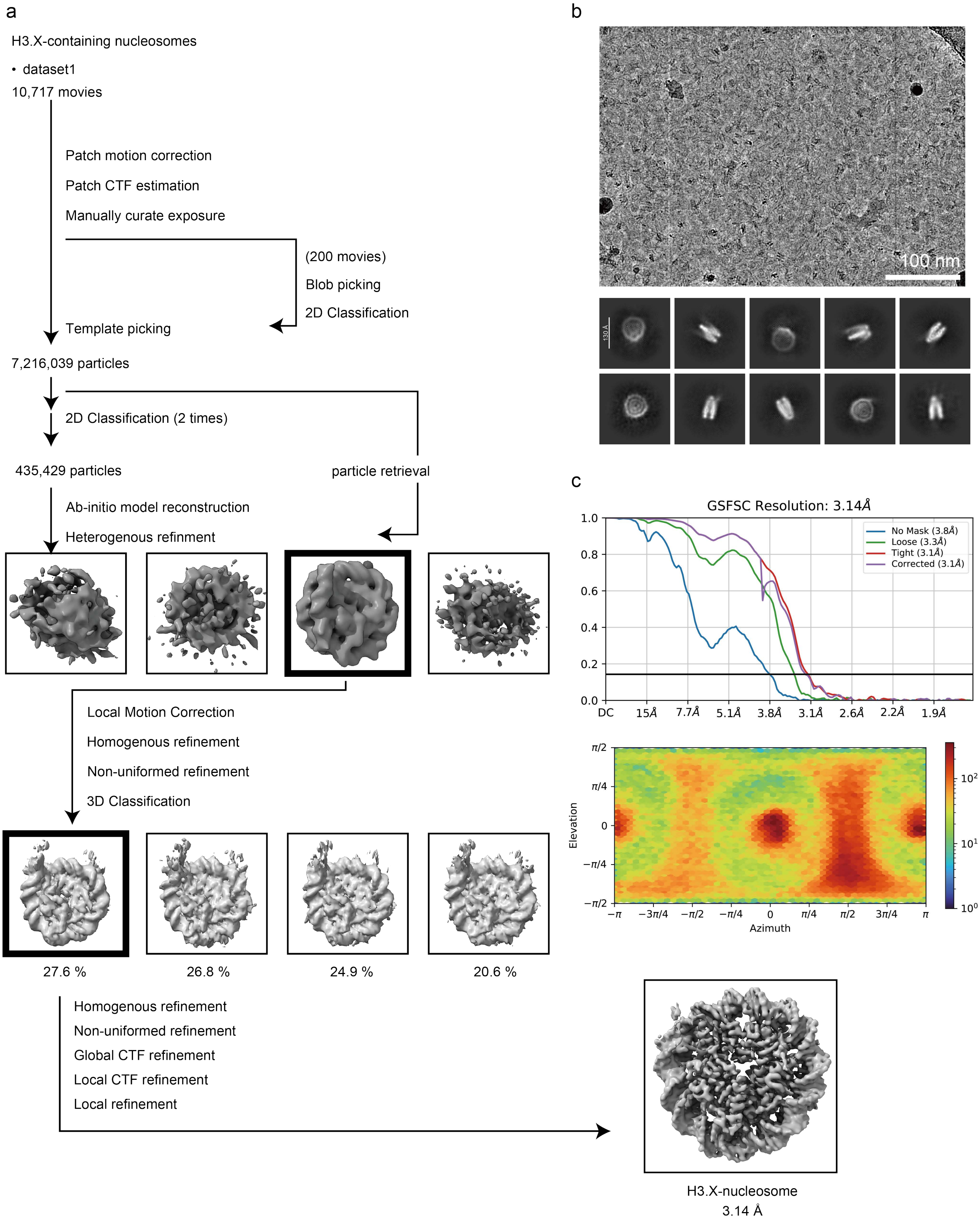


**Supplementary Figure 5. Data processing of H3.X-containing nucleosomes.**
**a,** Cryo-EM data processing workflow for H3.X-containing nucleosomes. **b,** Representative micrograph and 2D class images. **c,** Global resolution assessment by FSC curves at the 0.143 criteria (top) and the angular distribution of particle representation by viewing direction distribution plot (bottom).

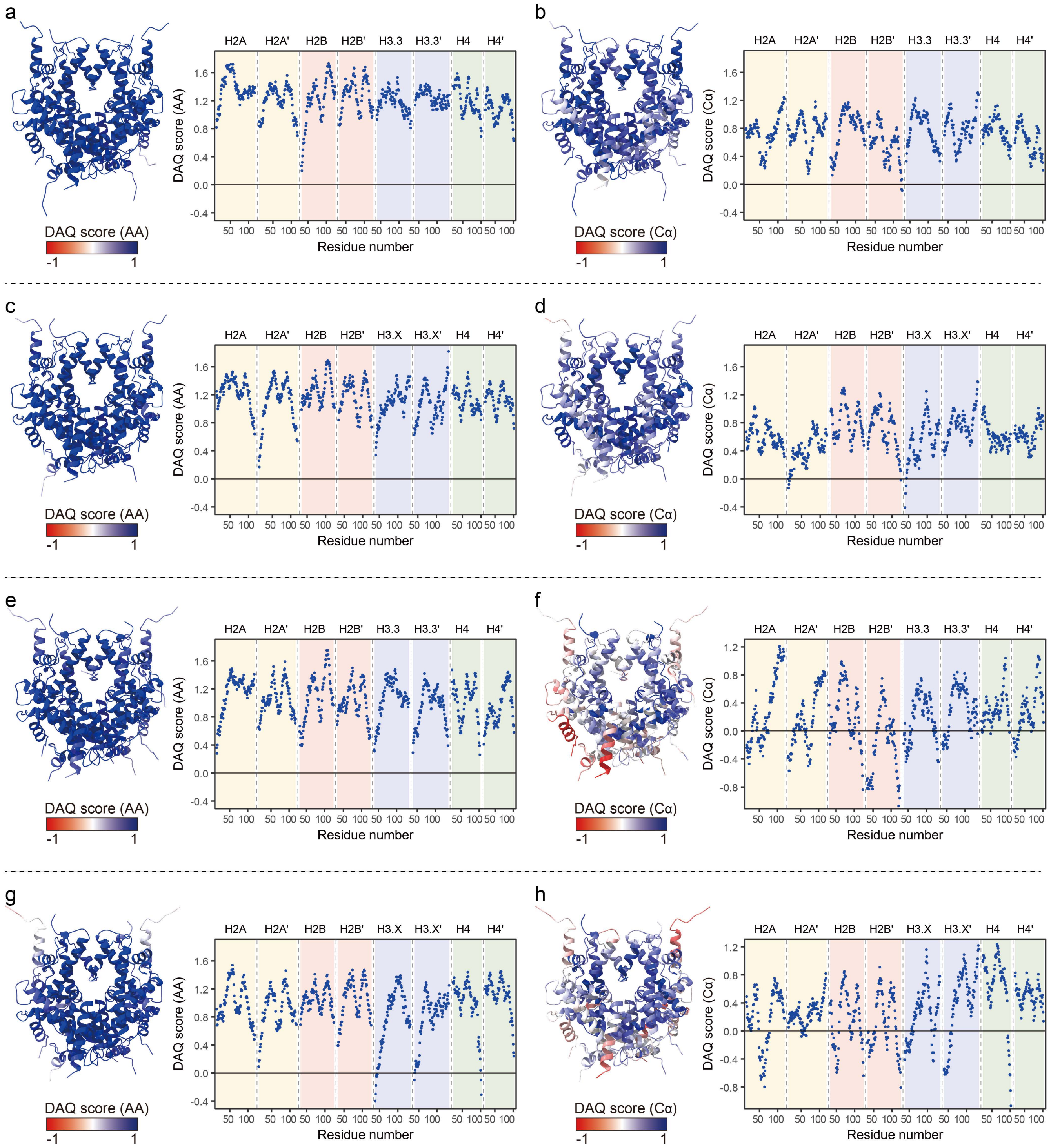


**Supplementary Figure 6. Structure validation of H3.3- and H3.X-containing nucleosomes using DAQ scores.**

**a-d,** Histone octamer colored according to side-chain-wise (**a**,**c**) and backbone-wise (**b**,**d**) DAQ scores of H3.3- (**a**,**b**) and H3.X-nucleosomes (**c**,**d**). The color code indicates DAQ values ranging from -1 (poor) to 1 (excellent); the accompanying dot plots show the estimated DAQ score at each residue. **e-h,** Cross-validation using side-chain-wise (**e**,**g**) and backbone-wise (**f**,**h**) DAQ scores of our H3.3-nucleosome map against the model of 3AV2 (**e**,**f**) and our H3.X-nucleoosme map against the model of 5AY8 (**g**,**h**).

a

H3.X
ARTKQTARKATAWQAPRKPLATKAARKRASPTGGIKKPHRYKPGTLALREIRKYQKSTQLLLRKLPFQRL
VREIAQAISPDLRFQSAAIGALQEASEAYLVQLFEDTNLCAIHARRVTIMPRDMQLARRLRGEGAGEPTLLGNLAL

b

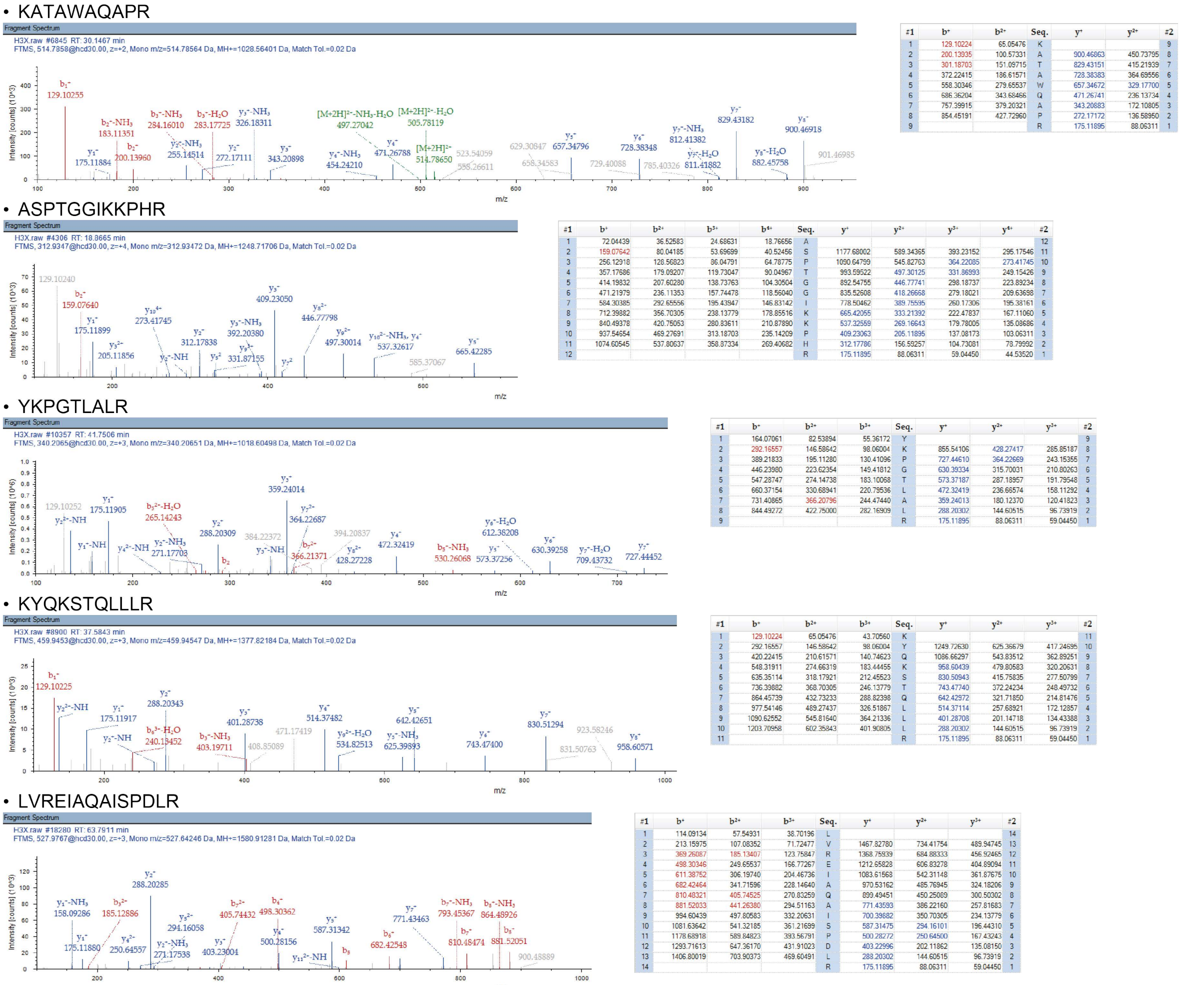


c

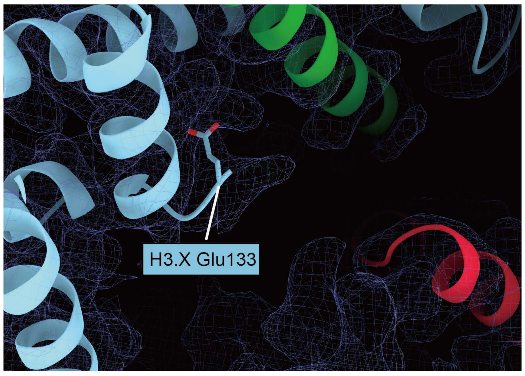


**Supplementary Figure 7. LC-MS/MS analysis of histone H3.X and C-terminal region is missing in the 3D structure.**

**a,** Peptide coverage of H3.X identified by LC-MS/MS analysis. Detected peptides by LC-MS/MS are highlighted in green. **b,** Fragment spectra of representative detected peptides. Detected fragmented ions are highlighted by blue or red in the right tables. **c,** Close-up view of the extended C-terminal region of H3.X in the nucleosomes.
